# The South American MicroBiome Archive (saMBA): Enriching the healthy microbiome concept by evaluating uniqueness and biodiversity of neglected populations

**DOI:** 10.1101/2025.04.03.647034

**Authors:** Benjamin Valderrama, Paulina Calderon-Romero, Thomaz F.S. Bastiaanssen, Aonghus Lavelle, Gerard Clarke, John F Cryan

## Abstract

The composition and function of the human gut microbiome has been linked to multiple health outcomes across all world regions, often with region-specific associations. Unfortunately, the extent to which microbiomes from different populations are characterised is limited by their economic resources. Over 70% of the sequenced human microbiomes come from analyses of European and North American populations, skewing our understanding by focusing excessively on just 15% of the global population. Thus, entire continents rely on results from research conducted in wealthier countries whose main findings are unlikely to generalize across other world regions. Moreover, statistical models perform poorly when applied to minorities, a blind spot with serious consequences in biomedicine, and which can only be addressed by analysing microbiome data from currently neglected areas. To address this problem, we created saMBA— the largest archive of gut microbiomes from South America, one of the world’s most biodiverse regions in terms of the gut microbiome of its inhabitants, yet the one with the fewest samples. ‘saMBA’ includes 33 gut microbiome studies, ∼73% of which were incorporated in a microbiome archive for the first time. By leveraging this resource, we uncovered a high biodiversity within —and uniqueness between— gut microbiomes across the continent, expanding the concept of the healthy microbiome to be more globally representative. Additionally, our results highlight that the gut microbiome biodiversity of this region remains far from fully characterized. We demonstrate how saMBA can guide new sampling efforts to better capture this diversity. Finally, the code deployed to build saMBA is compatible with that of a previous global compendium and is openly available to researchers from other underrepresented regions, fostering the inclusion of other neglected populations to accelerate microbiome research globally.

## Introduction

The microorganisms living in the human gut, collectively known as the gut microbiome, have been linked to multiple aspects of human health ^1,2^. Its composition is influenced by host genetics, and by environmental factors such as diet, lifestyle, exposure to xenobiotics, among others ^3–5^. Although initially the role of nature or nurture in shaping human gut microbiome composition was debated ^6^, it is currently accepted that while environmental factors are the main drivers, genetics make a comparatively smaller contribution ^5,7^. As geographic location represents an ensemble of genetic, environmental and cultural factors, world-region specific gut microbial signatures have been identified ^8^. Complementary observations on migrant individuals highlight the dynamic nature of their gut microbiomes, which tends to blend with the gut microbiomes of locals in a time-dependant manner ^9,10^. These observations indicate that healthy microbiomes and those associated to diseases are different among world regions, questioning the universality of microbiome-derived biomarkers of disease and indicating the probable need for more precise microbiome-based therapeutics. Indeed, recent efforts to define the healthy microbiome had advocated for a more even characterization of the gut microbiomes across world regions ^1^.

Notably, the extent to which microbiomes from different populations are characterised is limited by their economic resources. Nowadays, more than 70% of the public human microbiome data with known origin is from people in Europe or North America, despite that their combined population represents less than 15% of the global population ^11^. Consequently, entire continents are underrepresented, relying on results from research conducted in wealthier countries that may not face the same diseases that poorer countries struggle with ^12^, and whose main findings may fail to be generalized to other populations ^13^. For instance, in South America almost every country was identified as being underrepresented in a global-scale analysis ^11^. Thus, some local ^14,15^ and continent-wide initiatives are trying to increase the representation by taking more samples ^16^.

In addition to increasing the sampling effort in underrepresented communities, systematically evaluating publicly available microbiome data under a unified framework is a key front to advance our knowledge about these gut microbiomes. The integrated analysis of data from different studies can facilitate the discovery of patterns that may remain elusive when studying single cohorts. In that respect, the largest compendium of human gut microbiome to date, the Human Microbiome Compendium (HMC), was created by gathering 168,000 samples from all over the world ^8^. The HMC represents a big step forward in current-knowledge systematization. However, due to the scale of this analysis, authors only included projects where more than 50 samples were sequenced ^8^. Establishing such thresholds is a reasonable pragmatic decision considering the computational resources required for such analyses. However, it unintentionally impacts world regions in a way that exacerbates existing differences in sampling efforts, as poorer regions may have higher proportion of projects with less than 50 sequenced samples.

To address this limitation, we screened the international nucleotide sequence database collaboration (INSDC)^17^ and manually curated a list of 33 gut microbiome studies from South America, the world region with the fewest microbiome samples but one of the highest gut microbiome diversities among its inhabitants ^8^. Since the workflow used to build the HMC can’t be used by third parties, we recreated their workflow to analyse these previously excluded projects. Consequently, we used the same software used in the HMC, and followed the same quality criteria to filter projects, samples and sequences, thus facilitating the comparison between resources. Moreover, the workflow used to build the South American MicroBiome Archive (saMBA) is available on GitHub (https://github.com/Benjamin-Valderrama/saMBA-pipeline) and can be accessed by other researchers globally. We hope this facilitates the replication of our results and the inclusion of other neglected world regions thus accelerating microbiome research globally.

## Methods

### Identification of bioprojects included in saMBA

The European Nucleotide Archive (ENA) was used as an interface to systematically search the INSDC. Two authors (BV and PCR) screened the databases independently to identify bioprojects including samples from the gut microbiome of South American individuals. For each South American country, the following search term was used: “(microbiome OR microbiota) AND South American country”. These are two examples of terms used: “(microbiome OR microbiota) AND (Chile OR chile)” and “(microbiome OR microbiota) AND (Brazil OR brazil)”. Broad search terms without restriction of publication year were used to avoid early exclusion of potentially relevant bioprojects. Each bioproject from the list of results was then screened to assess its relevance based on the project description and associated published article. The final list of projects included in saMBA is available in Supplementary Table 1.

### Sample selection

Further selection of samples within bioprojects was required for reasons including: (1) some bioprojects included human samples along with environmental samples, or samples from other body sites, (2) bioprojects included samples with different sequencing strategies, like amplicon sequenced and whole genome sequenced samples, or (3) bioprojects included samples from humans living in other world regions. Thus, samples included in saMBA were limited to those flagged with the “amplicon” library strategy and that can be linked to human hosts living in any South American country at the time sampling took place, either by metadata on the original publication, or by metadata in INSDC.

Additionally, samples sequenced with technologies other than Illumina sequencing, such as MinION and 454 pyrosequencing were further excluded. This decision was supported based on observations made by the authors of DADA2, who suggest using different parameters to analyse pyrosequenced samples ^18^, and because Illumina was the most common technology used among the screened bioprojects. Notably, it was observed before ^8^ that the sequencing instrument “454 GS” is the default value for the “instrument” field when samples analysed with Mothur ^19^ are uploaded to INSDC. Therefore, a manual review of the original publications associated with each bioproject was required to determine the actual sequencing technology used. If the information in the research article did not match the information in INSDC, the information provided by authors in the article prevailed. A list of all samples used and the associated bioproject is available in Zenodo (https://zenodo.org/records/15050380).

### Analysis of individual projects included in saMBA

This workflow was built following that used in the creation of the HMC ^8^, as detailed in the methods section and the archived code shared by the authors (https://zenodo.org/records/13733642). This includes all steps in the analysis of individual projects, and the criteria to assess the quality of the projects, samples and ASV sequences included. The only deviation in those guidelines was that, to build saMBA, FASTQ files from each included bioproject were downloaded using fastq-dl v2.0.4. The software, versions and code deployed to build saMBA are publicly available on GitHub (https://github.com/Benjamin-Valderrama/saMBA-pipeline). Briefly, all identified projects were analysed, regardless of the number of samples sequenced. The library layout of the bioprojects (either paired- or single-end) was determined automatically by the saMBA workflow and analysed accordingly. The analysis of samples was performed in R v4.3.3 using the DADA2 ^18^ v1.30.0 package. After the analysis, the quality of the project was assessed using the same two criteria applied by the authors of the HMC ^8^ : First, the number of non-chimeric sequences over the total number of sequences. Projects were discarded if the percentage of non-chimeric sequences is <50% of input sequences. Second, projects with 5 or more of the first 10 samples with >25% of chimeric sequences were also discarded. Paired-end sequenced projects failing any of the two criteria were reanalysed as single end by discarding the reverse reads, as described in the HMC ^8^. Re-analysed projects not meeting the inclusion criteria were finally discarded. Resulting ASV-level count tables of each project were clustered at the genus level, generating two main outputs for each project successfully analysed.

### Building saMBA

Genus-level count table of each bioproject analysed were consolidated into one archive-wide table. Then, individual samples were examined and quality-filtered, as reported before in the HMC ^8^. Briefly, samples with less than 10,000 non-chimeric reads were discarded. Then, taxa with less than 1,000 reads across samples and taxa present in less than 100 samples were removed. Notice that although the HMC remove taxa present in less than 1,000 samples, the total number of samples in saMBA is less than in HMC, thus making this criterion more stringent in saMBA. After removing taxa with low prevalence, samples with less than 10,000 remaining reads were discarded. Finally, samples with more than 10% of the reads assigned to Archaeal taxa, or with more than 10% of reads unclassified at the phylum level were also discarded. Samples meeting the quality criteria across all included bioprojects were then consolidated into two count tables, one with ASVs and other at the genus level, configurating the main outputs of saMBA. The consolidated count tables before and after filtering low quality samples and taxa can be accessed online through Zenodo (https://zenodo.org/records/15050380). The GitHub repository with the workflow also contains the code used to consolidate and to filter low quality samples.

### Analysing saMBA

The consolidated and filtered genus-level table was then used in all analysis besides those performed to compare saMBA with the HMC. Analyses were performed in R v4.3.3. To estimate microbiome diversity, the Chao1 and Shannon alpha diversity indices were calculated using the package microbiome v1.24.0. To estimate the per-country microbiome uniqueness, distances between pairs of samples within each country were calculated using the Jaccard dissimilarity as implemented in the function vegdist from the package vegan v2.6.6.1. Figures depicting geographic data were generated using the package spData v2.3.3. Other figures were generated with ggplot2 v3.5.1 and patchwork v1.3.0. The scripts deployed to analyse saMBA and to generate the figures of this article are publicly available in an independent GitHub repository (https://github.com/Benjamin-Valderrama/saMBA-article).

### Local Representation Index

The calculation of the index was performed as described before ^11^. Briefly, global population data was obtained from the World Bank report for the year 2022, as it contains the most updated and trustable information we could find. That information was then used to calculate the country’s share of the total South American population. Another percentage was calculated for each country, indicating the country’s share of the total amount of samples included in saMBA. Then, we calculated the Local Representation Index (LRI): for countries with a sample percentage higher than its population percentage, we divided the former by the latter. In the opposite case, then the LRI was calculated as the negative reciprocal of this number. Thus, positive numbers represent how many times the number of samples is higher than what is expected based on country’s population share. On the other hand, negative numbers represent how many times the number of samples needs to be increased to match the expected number of samples from that country based on its share of the population. Notice that values calculated as described above were used to generate the relevant plots, which is different from the previously published approach ^11^, where the log10 transformation of the values was used for visualization.

### Subsampling simulations to estimate per-country and continental microbiome richness

To estimate how the current sampling effort allows to uncover regional microbiome richness, a custom subsampling simulation approach was used. First, counts were transformed into binary data, where 1 represents bacterial taxa present in the sample after applying the filters described above, and 0 was used otherwise. Then a custom function was used to perform a consecutive subsampling by randomly selecting an increasing number of samples and estimating the number of novel taxa identified across all samples (specific sampling depths are described on https://github.com/Benjamin-Valderrama/saMBA-article). This process was conducted for each country independently, and then for the entire continent. Importantly, the number of samples used in each iteration was below the total number of samples available for that country or the continent. This subsampling process was repeated 1,000 times for each depth, allowing to calculate the mean value and its standard deviation, which were later used in the final figures.

### Subsampling simulations to estimate microbiome richness of individuals from non-industrialized settings

When available, the manuscripts associated to the projects included in saMBA were screened to identify if the subjects from whom samples were taken lived in industrialised or non-industrialised settings. If the manuscript wasn’t available, information deposited on the INSDC was used. When a clear identification of the setting couldn’t be made, the project was not included in this analysis. Projects analysing samples from non-industrialised settings were further divided into 2 groups depending on whether the research topic included an examination of individuals suffering from a disease or diarrhoea, and studies characterising microbiomes of individuals without those conditions. For each group we performed a subsampling approach as described in the section above to estimate the richness revealed by current sampling efforts. The goal was to characterise the biodiversity of gut microbiomes in non-industrialized settings, which are generally deemed as more diverse. Additionally, it is recognized that people from non-industrialised settings are more expose to pathogens and parasites, which affect the composition of their gut microbiome, hence the subdivision between studies conducted on individuals with and without the conditions mentioned above.

### Comparing saMBA and the Human Microbiome Compendium

It has been reported that using different workflows to analyse the same biological sample can modify the interpretations of the data ^20^. To validate compatibility between saMBA and the Human Microbiome Compendium (HMC), we evaluated the similarity of the results given by both workflows when analysing the same samples. Note that the available HMC count table is from before the preprocessing steps described in the ‘Building saMBA’ section. Therefore, to ensure fair comparisons, the unprocessed saMBA count table at the genus level was used when comparing saMBA and HMC (Figure 2A and Supplementary Figure 3). All other results were generated using the filtered (processed) saMBA count table. Using R v4.3.3, the 7 most abundant phyla across all samples from South American subjects included in the HMC v1.0.1 were identified, and their relative abundances were determined for each sample analysed on both workflows. This allowed a qualitative assessment on the degree of similarity of the same sample analysed with the two workflows. Then, a quantitative assessment was sought. This time, Bray-Curtis dissimilarities were calculated using the R function vegdist from the package vegan v2.6.6.1 for each pair of samples analysed with both workflows. Dimensionality reduction was performed using the R function prcomp.

## Results

### Introducing the South American Microbiome Archive (saMBA)

The HMC ^8^, a previous microbiome archive, was built by performing an automated search of gut microbiome samples from humans across the different world regions. Additionally, note that projects including less than 50 samples were excluded from the HMC. A re-analysis of the HMC resource shows clear differences in the gut microbiomes of South American populations when compared to populations from Latin American and the Caribbean (Supplementary figure 1), which suggests that explorations at a finer geographic resolution are needed. Thus, we set to expand the catalogue of human microbiome samples from South American populations. After screening the INSDC, a manual curation of bioprojects with gut microbiome samples from South American populations (see methods), allowed the identification of 33 studies. The unified analysis of these studies (Supplementary figure 2) enabled the creation of the largest archive of gut microbiome from South American populations: saMBA. Noteworthy, ∼73% the studies were included in any archive for the first time. Interestingly, 30% of the studies included had less than 50 samples (Figure 1B), suggesting that the exclusion criteria used in building the HMC increases the underrepresentation of South America, and potentially, the global south. Studies included in saMBA contain microbiome samples from people from 9 out of 13 South American countries (Figure 1A and 1C), amounting to a total of 2,971 samples in the final output (Supplementary figure 2), with a mean of 110 ± 164 samples per study, and a median of 3.37×10^4^ of non-chimeric reads per sample (Figure 1D). Thus, saMBA represents the most comprehensive resource of gut microbiomes from South Americans.

**Figure 1:**
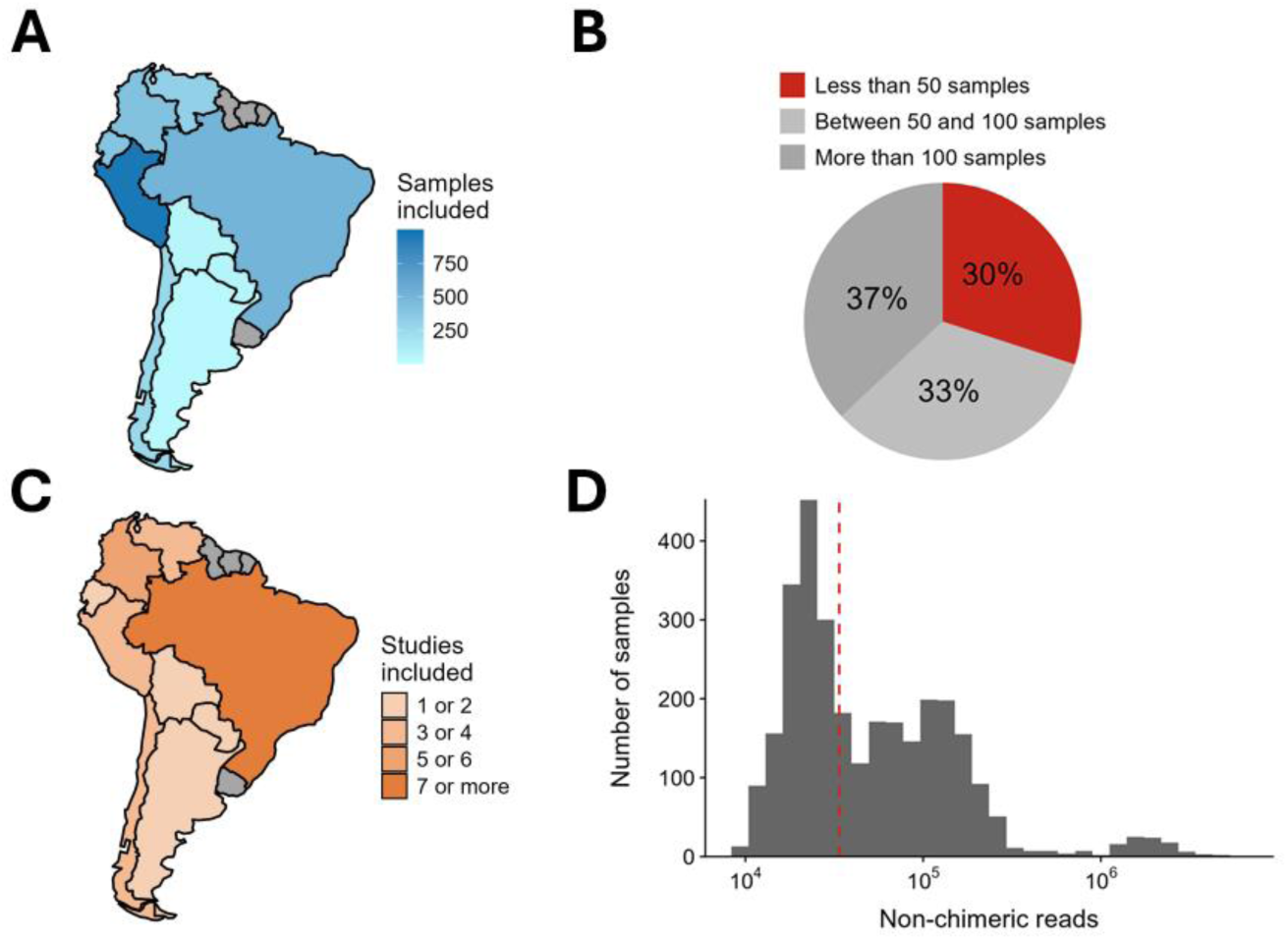
Overview of saMBA’s width across the South American continent. (A) Number of samples included in saMBA from each South American country. Darker blues indicate higher number of samples, grey indicates countries without data (B) Percentage of studies including less than 50 samples (red), between 50 and 100 samples (light grey), and more than 100 samples (dark grey) across countries. (C) Number of projects included in saMBA from each South American country. Darker oranges indicate higher number of projects, grey indicates countries without data. (D) Number of non-chimeric reads detected in samples across all projects included in saMBA. Red dotted line represents the median of the distribution.

### saMBA unveils high biodiversity across the continent

The manual curation of studies included in saMBA allowed the addition of 24 studies not included in previous archives. This highlights the need for manual curation of studies metadata for more accurate assessments of regional microbiomes. By leveraging saMBA, we identify a total of 2,246 genera across samples, more than doubling the current number of genera known to be present in the gut microbiomes of the region ^8^. Indeed, 60% were not included as part of South American gut microbiomes in the HMC (Figure 2A), the largest archive including South American microbiomes. Interestingly, 876 of the 911 genera (∼96.2%) identified in the HMC were also detected in saMBA, which suggests compatibility between resources. A more extensive analysis of their compatibility is available as supplementary material (Supp. Figure 3).

**Figure 2:**
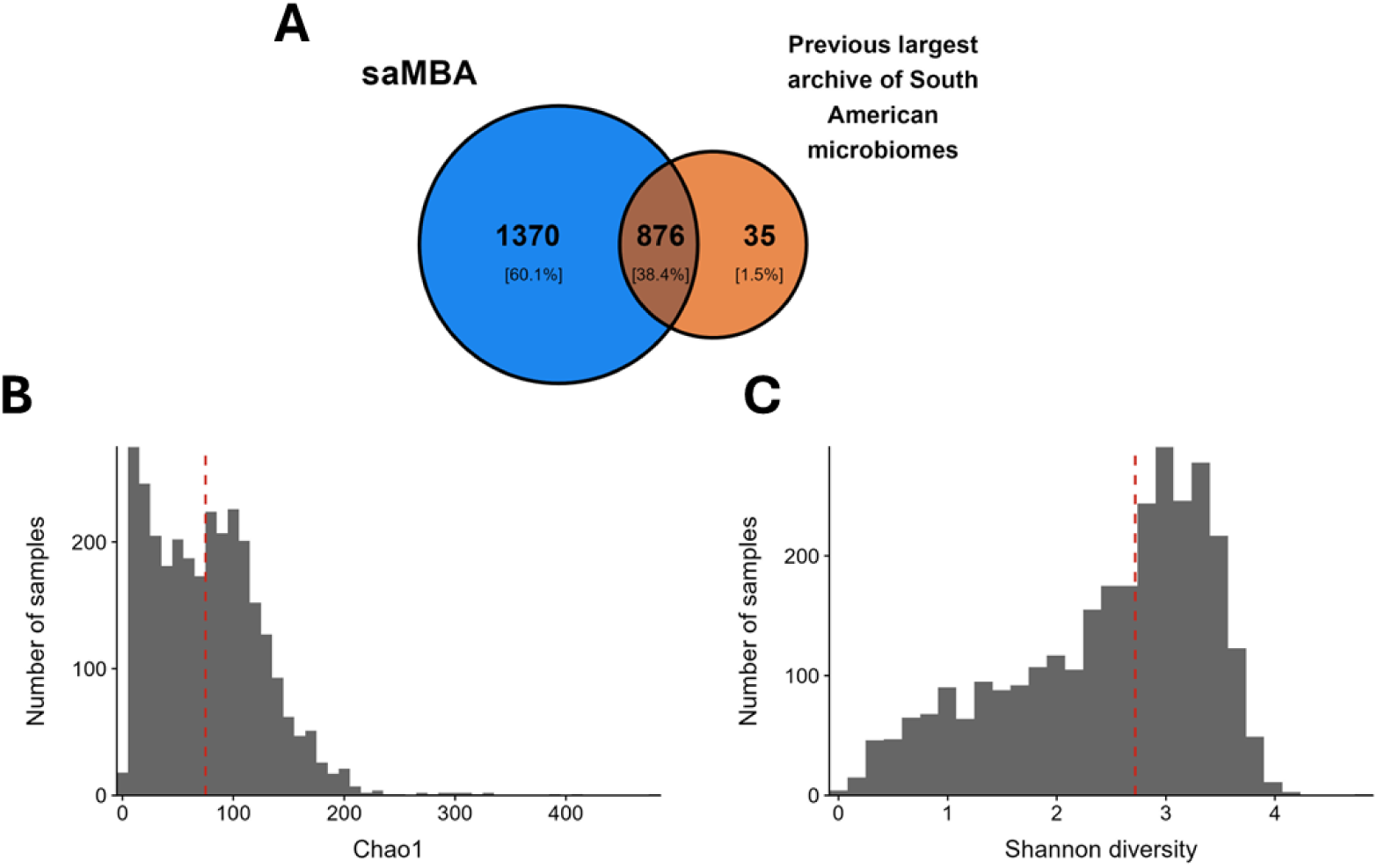
saMBA expands the number of identified bacterial taxa and shows a high degree of diversity across the continent. (A) Venn diagram showing that saMBA (in blue) covers most of the already identified genera in the region by a previous global compendium ^8^ (in orange) while more than doubles the current number of genera identified in the world region. (B) Distribution of Chao1 estimates of alpha diversity across all samples included in saMBA. Red dotted line shows the median value. (C) Distribution of Shannon estimates of alpha diversity across all samples included in saMBA. Red dotted line shows the median value.

Additionally, saMBA unveils a high biodiversity within the continent, as shown by two alpha diversity metrics (Figure 2B and 2C). The median of the Chao1 and Shannon indices for the continent were 85 and 2.77 respectively, which suggests a high number of different genera (Chao1) and a tendency to a more even distribution of them (Shannon) in the gut microbiomes of South American individuals.

### Biodiversity across the continent is most likely underestimated: New sampling efforts must account for the biodiversity and uniqueness of each country

We hypothesized that due to the inclusion of new samples from countries not included in previous archives, new estimations of the total biodiversity across the country would be more accurate. Our subsampling simulation analysis (see methods) showed that, although some countries are close to reach a plateau, none has done so yet (Figure 3A), suggesting that newer samples taken in most South American country will likely identify new genera not yet included in saMBA or any other compendium.

**Figure 3:**
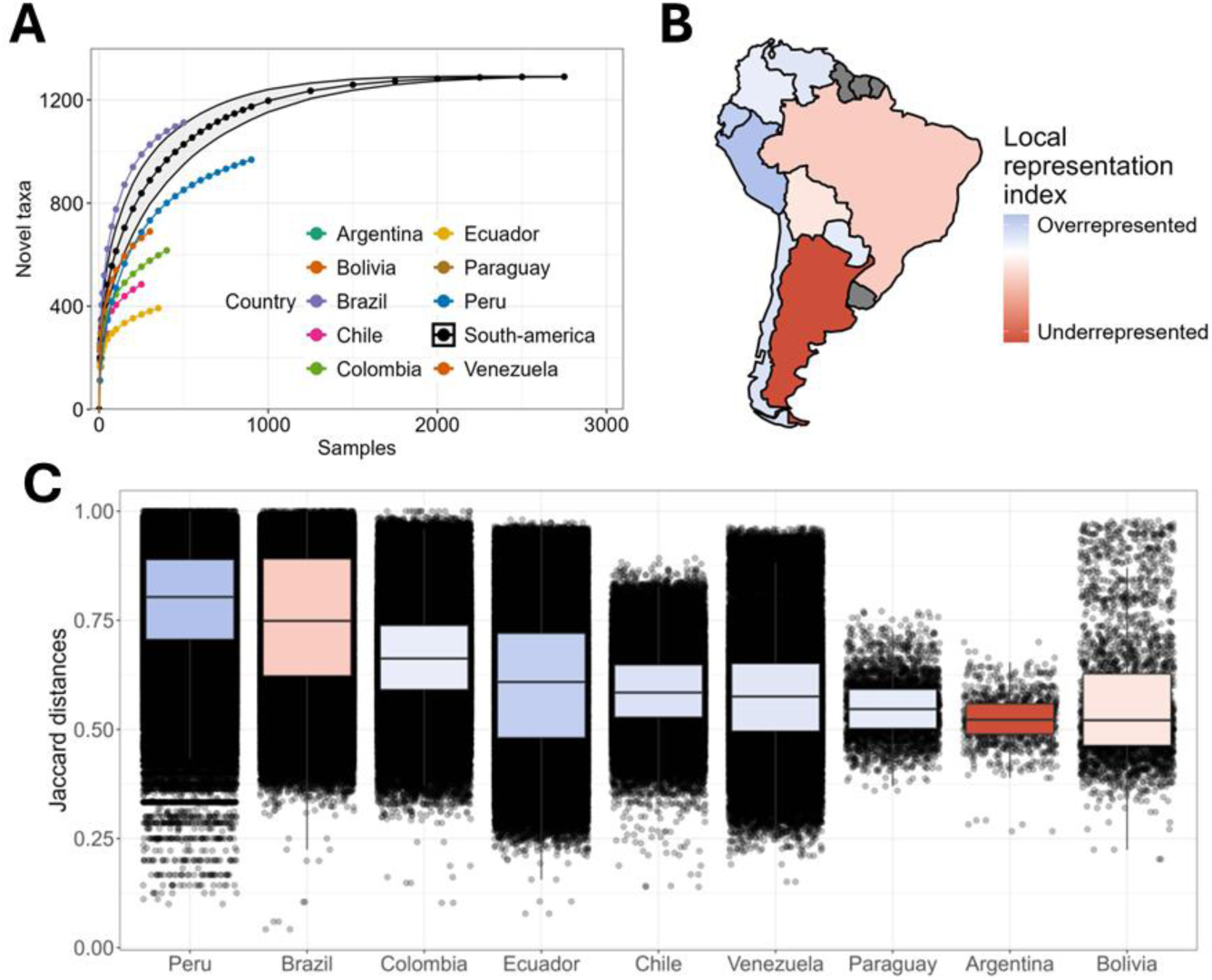
Biodiversity across the continent is likely underestimated, and future sampling efforts should consider biodiversity and uniqueness within each country. (A) Novel taxa identified when subsampling an increasing number of samples from each country. Each dot represents the mean value of 1,000 calculations of the number of novel taxa identified when subsampling at varying depths (i.e., number of samples). Each colour is a country. Black represents the estimates for the entire continent. They grey area around the continental estimate (black line) represents the SD. (B) Continental map of the Local Representation Index (LRI). Shades of blue represent countries where the proportion of microbiome samples is bigger than its share of the continental population, whereas shades of orange show the opposite. Grey indicates countries without available data. (C) Boxplot and distribution of all Jaccard distances calculated using every pair of samples available for each South American country. The colours of each boxplot match the colour of the country given by its LRI, as represented in panel B. Countries are organized from left to right in decreasing order based on median Jaccard distance.

Next, we repeated the analysis clustering all samples at the continental level (Fig 3A, black line). Our results show that the continent reached a plateau when subsampling at a depth of 2,000 samples (∼67% out of the total samples available in saMBA). Considering that most countries haven’t plateaued yet (Figure 3A), this may suggest that the gut microbiome biodiversity in the continent is still underestimated. This is further implied by our subsampling analysis conducted on projects studying subjects living in non-industrialised settings like the amazonian communities (Supplementary Figure 4), showing a yet unveiled biodiversity in their gut microbiomes. Thus, more samples are required to accurately characterise the regional biodiversity. However, we anticipated that sampling from different countries will have varying impacts in our understanding of South American gut microbiomes.

Next, we aimed to identify countries that can maximize the chances to identify yet unobserved bacterial taxa when collecting and analysing future samples. Thus, we first calculated a Local Representation Index (see Methods). Our results identified 6 overrepresented countries where the proportion of gut microbiome samples is higher than expected based on their share of the population, and 3 underrepresented countries, characterised by the opposite (Figure 3B). Then, for each country, the Jaccard distance between every pair of samples was calculated (Figure 3C). Higher Jaccard distances between samples indicate lower number of shared genera. Thus, countries with higher median Jaccard distances indicate a higher level of gut microbiome uniqueness within their population. Our results imply that future samplings should consider how well represented each country currently is, and the uniqueness of the gut microbiomes of people living in those countries. Considering both aspects to inform where to conduct newer sampling efforts can help maximize the return on investment, which is especially relevant in regions where resources are more limited.

## Discussion

Here we introduced saMBA, the largest archive of gut microbiome samples from South American countries yet created (Figure 1), along with the code required to reproduce the analysis of this and other neglected global regions. Our results show a significant expansion in the number of genera found in samples across this world region, more than doubling what a previously released global compendium reported ^8^ (Figure 2A). Additionally, by leveraging saMBA, we generated the most accurate assessment of regional biodiversity to date and provided guidelines on where new sampling efforts are most needed—an analysis that, to the best of our knowledge, is the first of its kind.

Our approach illustrates how semi-automated screening of databases (as performed in the creation of saMBA) can improve our sensibility to include relevant datasets (Figure 1B) not detected by fully automated approaches. Nevertheless, semi-automated approaches are not always feasible, as they are time consuming, and they are more prone to error as the size of the search space increases. This further emphasise the relevance of defining small search spaces (such as just one continent) when creating newer compendiums aiming to expand previous global efforts. Attending to this reality, the code deployed in the creation of saMBA was made public (https://github.com/Benjamin-Valderrama/saMBA-pipeline), so researchers from other world regions can generate their own archives.

By leveraging saMBA, we identified a high degree of biodiversity within the gut microbiomes across the continent, as indicated by two alpha diversity indices: Chao1 (Figure 2B) and Shannon (Figure 2C). Indeed, the distribution of Shannon diversities in South America follows a similar shape to that observed in a previous global compendium ^8^ but it’s shifted to the right, suggesting that people in this continent tend to harbour more diverse gut microbiomes than people living in other regions. The previous global compendium didn’t provide a distribution of Chao1 indices, making it impossible to compare. However, it is interesting to note that while in some samples more than 150 different genera were identified, others show less than 50 (Figure 2B). A closer inspection identified the articles where those samples were taken. Most samples in the high end of the distribution come from one early work characterising the gut microbiome from adult individuals living in the Venezuelan amazon ^21^. Interestingly, one of the main conclusions of that work is that those gut microbiomes are much more diverse than those of people living in industrialized environments. On the other hand, most samples in the lower end of the distribution were traced back to two studies. One conducted in infants with intestinal inflammation ^22^, and the other in undernourished children with diarrhoea ^23^. It is notable that results from individual research articles were replicated during this broader analysis, validating the parameters selected for the bioinformatics analyses used to build saMBA.

Research conducted in individual South American countries has consistently pointed out the high diversity of the gut microbiome of their inhabitants ^21^. However, accurate estimations of the true extent of this biodiversity were missing, probably due to the narrower scale of previous analyses, which focused on single populations rather than across the continent. Thus, our assessment of the regional biodiversity is the first of its kind and extent. By applying a subsampling simulation approach (see methods section), we showed that more sampling efforts are still required to accurately characterising all microbiome genera present in the gut microbiome of people living in South America (Figure 3A). Thus, while it is well accepted that the gut microbiomes of South Americans are highly diverse, we are probably still underestimating its real biodiversity. Interestingly, previous results have shown that Latin America and the Caribbean (which encompasses South America) is the world region with the lowest number of gut microbiome samples ^8^. Together, these results emphasize the need for more sampling across the continent, a challenge which new local scientific initiatives have already started to tackle ^16^.

However, the question of where those samples should be taken is not trivial, especially in regions where the scientific budget is scarce. Thus, we determined a Local Representation Index for each country (Figure 3B). We aimed to assess if the proportion of microbiome samples from each country matches the proportion of the South American population living there. This allowed us to identify underrepresented countries in the region, an essential guide to where future sampling efforts should be conducted. Additionally, the uniqueness of gut microbiomes within each South American country (Figure 3C) must also be considered. In this regard, we identified countries having the largest internal heterogeneity, which may indicate countries with highest chances to identify yet unobserved bacterial taxa when collecting and analysing future samples.

Although these results may help in deciding where to sample next, we recognize some limitations arising from the high levels of heterogeneity among South American countries. For instance, while Sao Paulo city has around 44 million habitants, a minor yet often studied part of their population lives in the Brazilian amazon region, with close to zero contact with the industrialized world. It’s expected that these populations will differ dramatically. Additionally, social inequities —that can shape our gut microbiomes in multiple ways— are prominent within South American countries like Chile ^24^. Regarding our own analysis, while we identify Argentina as being greatly underrepresented (Figure 3B), it also shows one of the lowest microbiome uniqueness (Figure 3C), suggesting that newer samples from the country may add little novelty. However, a closer inspection reveals that the three ^25–27^ included studies carried on Argentinian subjects sampled only individuals living in industrialized cities. Thus, sampling individuals living in other contexts may still bring valuable information. In contrast, Peru was identified as overrepresented (Figure 3B), but it also has highest level of heterogeneity (Figure 3C). This makes sense when considering that most samples come from individuals living in the amazon, which suggests that further bacterial biodiversity may remain to be discovered in the country. Thus, despite the progress this analysis represents, further finer scale analyses accounting for diverse country-specific realities must be done when deciding where to take future samples.

## Limitations

Although saMBA represents a major advance in understanding the gut microbiome with a global perspective, it has some limitations. First, saMBA focuses on amplicon data, specifically 16S, thus overlooking the mycobiome and virome. Future work should include whole genome sequencing data, allowing for the identification of other members of the gut microbiome, as well as a more detailed characterization of the bacteriome. Similarly, this work focusses on faecal samples. Importantly, the code used to build saMBA can be used to build archives of other body sites without further modifications. Additionally, the metadata associated with each sample heavily depends on what was made available by the original authors of each study, which in some cases can be scarce. Further improvements to this resource include allowing interested researchers who are submitting their study to INSDC, to share the accession codes to their samples and metadata with saMBA. Researchers interested in doing so are welcomed to open an issue on GitHub (https://github.com/Benjamin-Valderrama/saMBA-article).

## Conclusions

‘saMBA’ deepens our understanding of the different stable states of the human gut microbiome, thus expanding our understanding of a more globally representative microbiome. The workflow deployed to build saMBA was made available to other researchers and it is compatible with the HMC —a previously released global compendium. Thus, our work provides the impetus for the inclusion of other neglected populations to accelerate microbiome research globally, and guidance for newer sampling efforts taken place in South America.

## Supplementary material

### World regions used in previous work are too broad, justifying the analysis of gut microbiomes in narrower areas

The HMC ^8^, a previous work performed an automated search of gut microbiome samples from humans across the different world regions defined by the United Nations. Considering the diverse lifestyles within the ‘Latin America and the Caribbean’ region, we hypothesised that the gut microbiome of people living in Central and South America would be substantially different, potentially justifying narrower efforts to better characterise the subpopulations. A re-analysis of the HMC resource shows differences in the gut microbiomes of people in those subregions. While the abundance of Bacillota and Methanobacteriota are higher in subjects from Central America, South Americans show higher Bacteroidota and Pseudomonadota. This result suggests that explorations at a finer geographic resolution are justified.

**Supplementary Figure 1:**
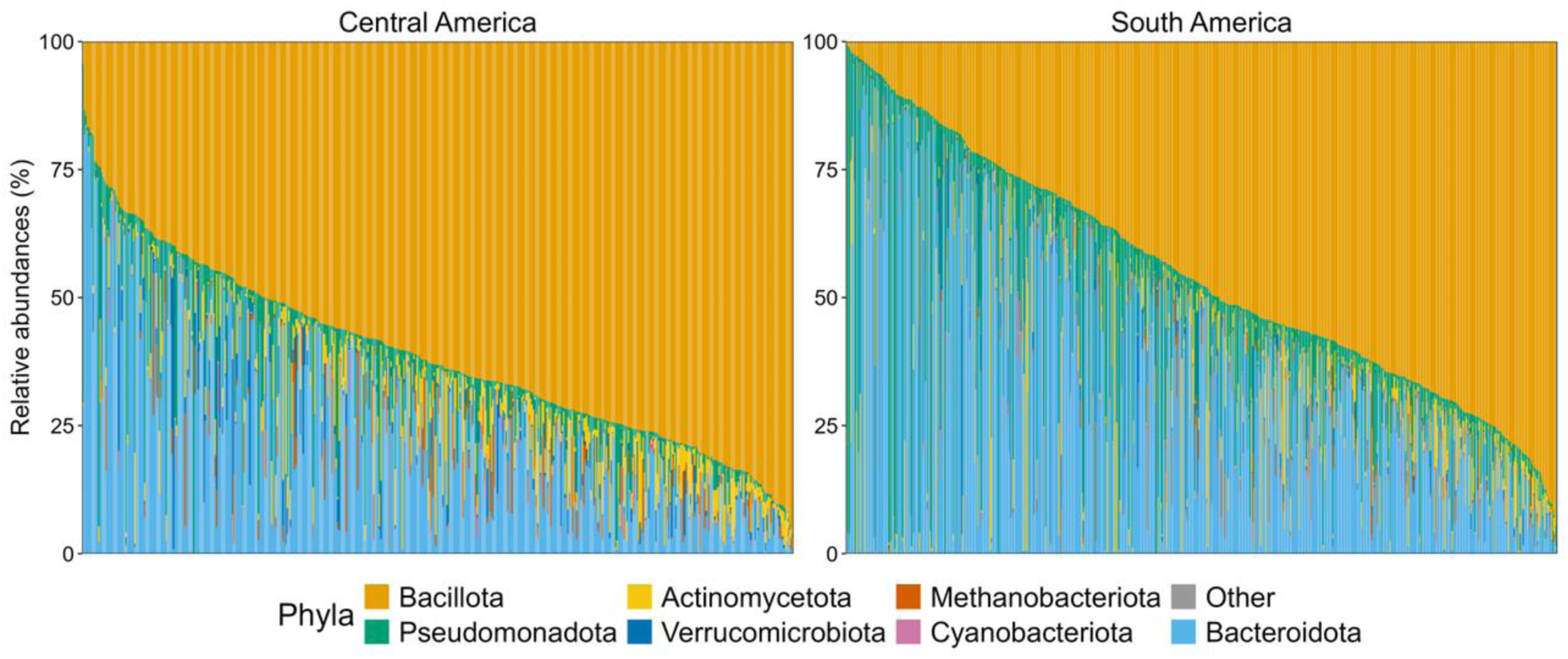
Microbiome samples from Central and South America show different compositions, justifying an independent exploration of each region. Panel shows the relative abundance of the most abundant phyla identified in Central America (left panel) and South America (right panel). Each colour depicts one of the 7 most abundant phyla across Latin America and the Carribean. Grey represents the relative abundance of all other phyla. Samples in each panel are ordered by their relative abundance of the phylum Bacillota.

### Schematic of the workflow used to build saMBA improves transparency in the bioinformatics analysis of samples

To improve transparency in our work, we show the workflow deployed in the analysis of each project included in saMBA. Each step was described in the methods and implemented as described in the GitHub repositories linked on each methods section. As a complement, the following schematic illustrates the steps of the workflow in a visual manner. Along with each step of the workflow (see the first column), we provide a small description of it (second column) and the number of samples successfully completing it (third column).

**Supplementary Figure 2:**
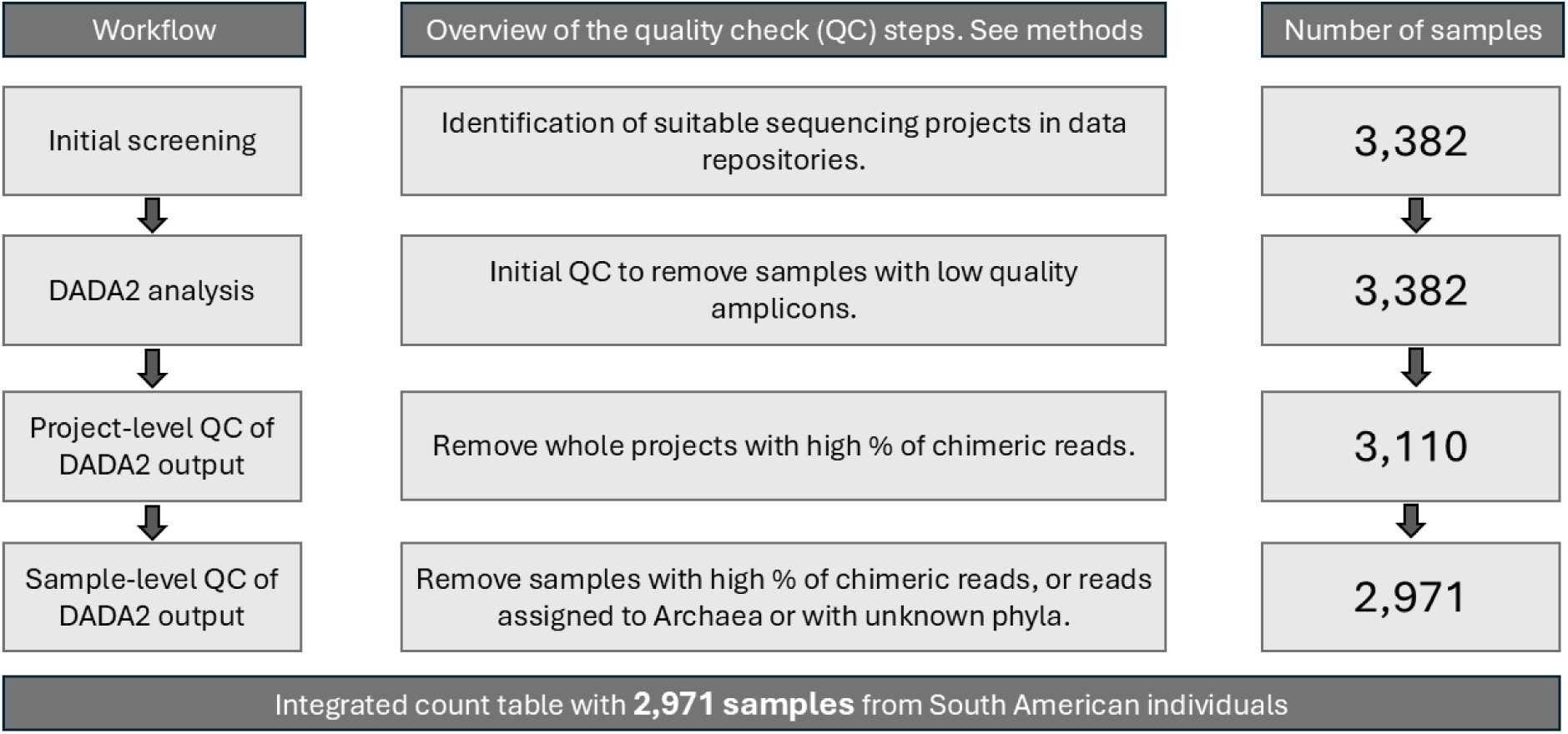
Number of samples successfully completing each step in the analysis workflow. Each step in the workflow has a brief explanation that matches the information available in the methods section.

### The community profiling of samples analysed with saMBA and the HMC workflows are concordant

It is known that different processing workflows can generate slightly different results when applied to analyse the same dataset. Since the workflow developed to build the HMC was not available for public reuse, we aim to recreate the workflow following the details provided by the authors in the method section and non-reusable code shared on an archived GitHub repository ^8^. The goal was to replicate the workflow as closely as possible. To examine how well we matched their processing steps, we followed qualitative and quantitative approaches. First, we compared the relative abundances of the seven most abundant phyla in samples included in both workflows (Supplementary Figure 3A). Although Actinomycetota was more abundant in some samples analysed with the saMBA workflow when compared to the same sample analysed with the HMC workflow, the overall relative abundances are concordant for most samples.

Next, we quantified the dissimilarity between samples analysed with each workflow to determine how different are the results generated by each when analysing the same sample. If the same sample is analysed with different workflow and they are highly dissimilar, it implies that the source of that dissimilarity is technical, as it comes from small differences in the bioinformatics analysis. However, if different samples are analysed with the same workflow and they are highly dissimilar, then the source is biological, not technical. Thus, one indication that workflow results are compatible is that the average dissimilarity resulting from analysing the same sample with different workflows is lower than the average dissimilarity of different samples analysed with the same workflow. In other words, if the biological sources of dissimilarity are on average higher than the technical sources (Supplementary Figure 3B).

To provide a reference, we added two other groups, where we calculated the dissimilarity of different samples analysed with different workflow (i.e., technical and biological source of dissimilarity), and the same sample analysed with the same workflow (i.e., no dissimilarity). Although some samples analysed with different workflows show high levels of dissimilarity, the boxplots show that more than 75% of the data in that group show less dissimilarity than any different samples analysed with the same workflow. Additionally, approximately 25% of the data shows no dissimilarity. This suggests that in most cases, the results generated by the workflow used to build the HMC and saMBA generate compatible results (Supplementary Figure 3B).

Finally, we sought to explore the overall dissimilarities between workflows (Supplementary Figure 3C). The ordination plot shows that although there is variability across samples in the first two dimensions, there is considerable overlap in the space used by samples analysed with different workflows. This is consistent with the results shown in Supplementary Figure 3B.

**Supplementary Figure 3:**
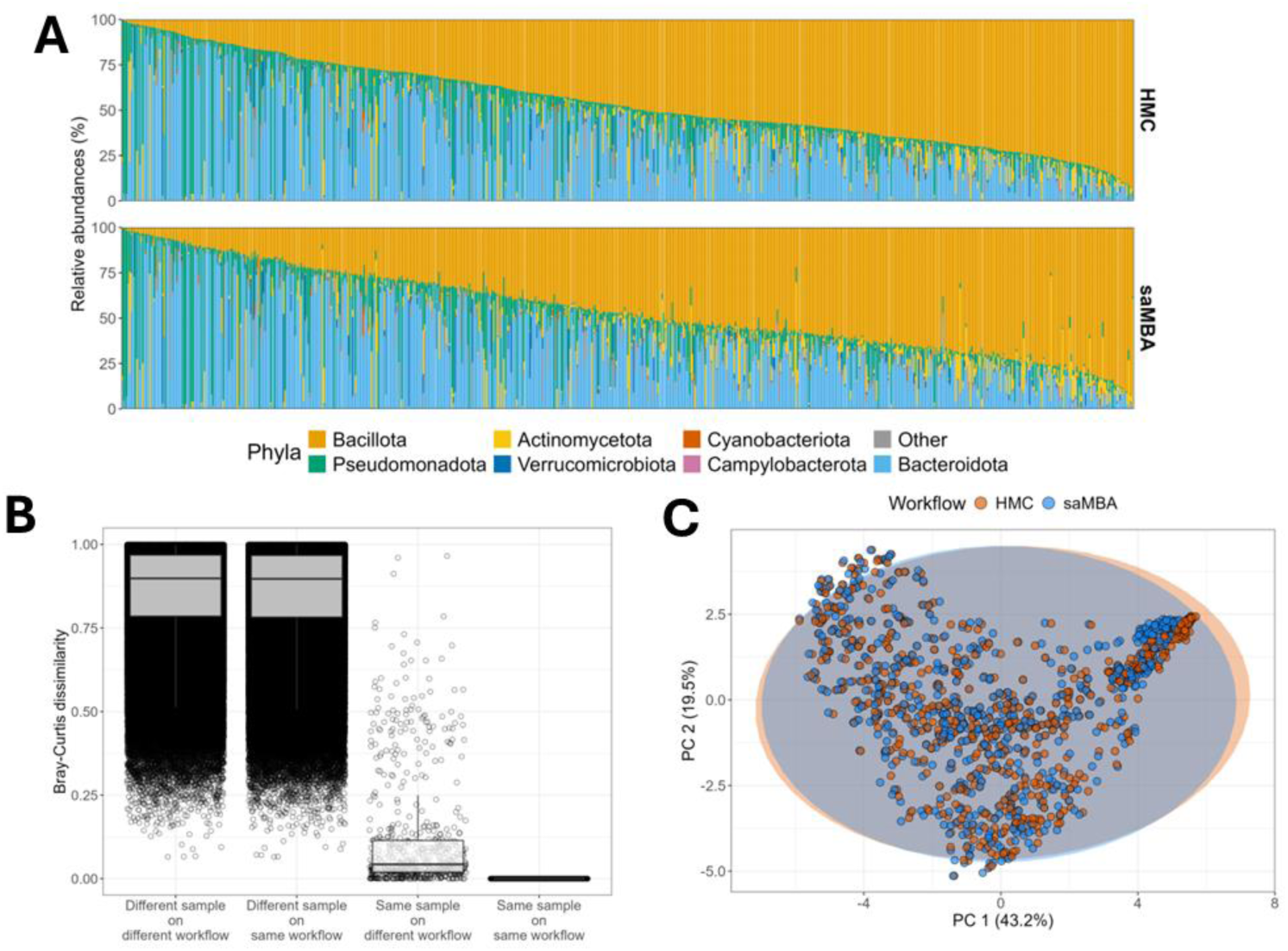
Qualitative and quantitative assessment of the dissimilarity between results generated by the HMC and saMBA workflows. (A) Relative abundances at the phylum-level in samples included in the HMC (top) and in saMBA (bottom). Each colour depicts one of the 7 most abundant phyla, and grey represents the relative abundance of all other phyla. (B) Boxplot of the Bray-Curtis dissimilarities between pair of samples. Pairs are either (1) different samples analysed with different workflows, (2) different samples analysed with the same workflow, (3) the same sample analysed with different workflows and (4) same sample analysed with the same workflow. (C) PCA plot of using Bray-Curtis dissimilarities. Orange depicts samples analysed with the HMC workflow, and blue those analysed with the saMBA workflow.

### Non-diseased individuals living in non-industrialised settings across the region show a yet unveiled gut microbial biodiversity

We recognise the heterogeneity of lifestyles within South American countries. For instance, in Brazil, cities like Sao Paulo have around 44 million habitants, whereas its amazonian regions are inhabited by small groups of people with limited contact with the industrialized world. Thus, we conducted a subsampling simulation analysis (see methods) using seven projects analysing gut microbiomes of individuals living in non-industrialized settings. We further characterised projects based on whether they included individuals with infectious diseases or diarrhoea or not, completing a total of 1,072 and 446 samples, respectively. Our results (Supplementary Figure 4) suggest the existence of a yet unveiled gut microbiome biodiversity in South Americans living in non-industrialised contexts, specifically in those not suffering from diseases.

**Supplementary Figure 4:**
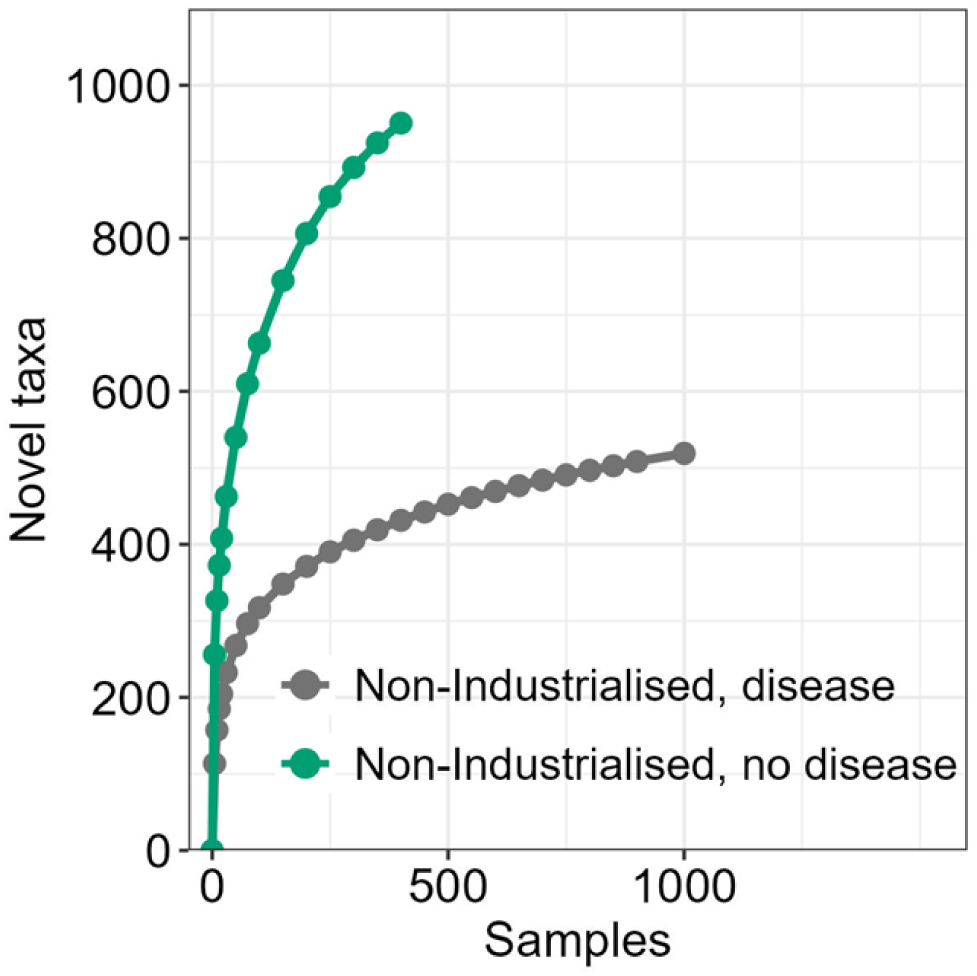
Gut microbiome biodiversity estimations in subjects living in non-industrialized regions. Novel taxa identified when subsampling at an increasing number of samples obtained from South Americans living in non-industrialised settings. Each dot represents the mean value of 1,000 calculations of the number of novel taxa identified when subsampling at varying depths (i.e., number of samples). Green represents the biodiversity estimates using samples taken from individuals living in non-industrialised settings without disease, whereas Grey represents the same for individuals living in non-industrialised settings with infectious disease or diarrhoea.

## Funding

APC Microbiome Ireland is funded by Research Ireland (previously Science Foundation Ireland, SFI grant code 12/RC/2273_P2).

## Conflict of interest

JFC has been an invited speaker at conferences organized by Bromotech, Yakult and Nestle and has received research funding from Nutricia, Dupont/IFF, and Nestle. GC has received honoraria from Janssen, Probi, Apsen, and Ingelhem Boehringer as an invited speaker; is in receipt of research funding from Pharmavite, Fonterra, Reckitt, Nestle and Tate and Lyle; and has been paid for consultancy work by Yakult, Zentiva, Bayer Healthcare and Heel Pharmaceuticals. This support neither influenced nor constrained the contents of this manuscript. All other authors declare no conflicts of interest.

## Acknowledgments

BV would like to thank Dr. Richard J. Abdill for the helpful discussions by e-mail about the implementation of the workflow used to build the Human Microbiome Compendium. BV thanks Dr. Samuel Miravet-Verde for the discussions about the automated screening of the INSDC.

## Author contributions

BV conceptualized the project together with PC-R, TFSB, AL, GC and JFC. BV and PC-R conducted the screening and identification of relevant bioprojects. BV curated the final list of bioprojects included. BV wrote the code to build saMBA. BV performed the data analysis. BV prepared the first draft. BV, PC-R, TFSB, AL, GC and JFC read and edited the manuscript. GC and JFC acquired the funding.

## Notes

https://zenodo.org/records/15050380

https://github.com/Benjamin-Valderrama/saMBA-pipeline

https://github.com/Benjamin-Valderrama/saMBA-article

